# *Drosophila* male genitalia rotation depends on permissive remodeling of the posterior abdomen

**DOI:** 10.64898/2026.03.27.714580

**Authors:** Nuria Prieto, David Foronda, Paloma Martín, Eléanor Simon, Marcus Bischoff, Stéphan Noselli, Ernesto Sánchez-Herrero

## Abstract

One of the most characteristic morphogenetic processes in *Drosophila* is the 360° rotation of the male pupal genital disc. This movement is driven by the myosin Myo1D, whose expression in the genital disc is controlled by the Hox gene *Abdominal-B*. The rotation takes place in contact and relative to the posterior abdomen, yet the contribution of abdominal tissues has remained unclear. Here we show that normal genital disc circumrotation requires active remodeling of posterior abdominal larval epidermal cells that contact the rotating terminalia. Preventing apoptosis in these cells, or increasing EGFR signaling, delays their extrusion and results in incomplete rotation without altering rotational chirality. In parallel, elimination of Extracellular Matrix by Metalloproteinase 1 in these cells, although without leading to their extrusion, is also strictly required for genital disc circumrotation. Inhibition of this metalloproteinase activity leads to persistence of collagen IV and incomplete rotation, revealing an independent requirement for Extracellular Matrix clearance at the disc–abdomen interface. By contrast, genetic conditions that prevent formation or elimination of the male A7 segment do not necessarily impair genital disc rotation, demonstrating that A7 suppression and circumrotation are separable processes. These findings identify posterior abdominal tissue remodeling as an essential extrinsic requirement that enables genital disc circumrotation.

## Introduction

Organ formation in bilaterians takes place in most cases symmetrically on the left and right sides. There are, however, many examples of left-right asymmetry, which determines different position, structure, or coiling of left and right organs (Coutelis et al., 2014). Studies in *Drosophila* identified the Myosin 1D protein as a key factor in determining asymmetries, and in all these cases the tissue twists or rotates in the same direction. A prominent example is the rotation of the male genital disc during pupal development.

The *Drosophila* male genital disc, located at the back of the larva, comprises three segments, A8, A9 and A10, which form the genitalia and analia (the terminalia) (Nöthiger et al., 1977). During early pupal development the disc everts, and about 24-25h after puparium formation (APF) it starts a 360° rotation that is always clockwise (when viewed from the posterior end of the animal) (Gleichauf, 1936; Speder et al., 2006; Hozumi et al., 2006; Suzanne et al., 2010; Kuranaga et al., 2011). This movement is driven by the polarized distribution of Myosin II and asymmetric cell intercalation (Sato et al., 2015), and its direction (dextral or sinistral) can be unambiguously scored by observing the looping of the spermiduct, which connects external and internal genitalia and turns around the hindgut. This coiling allows discriminating wild-type rotation from cases in which the genital disc fails to rotate, since the external arrangement of genitalia and analia in both cases is indistinguishable (Adám et al., 2003).

The start of rotation depends on the Hox gene *Abdominal-B* (*Abd-B*), (Coutelis et al., 2013), and on *myo1D*, which also sets the clockwise (or dextral) direction of rotation. In *myo1D* mutants the genital disc rotates but with inverted chirality (sinistral) (Speder et al., 2006; Hozumi et al., 2006), and in *Abd-B* mutants, where *myo1D* is absent, there is no rotation, either dextral or sinistral (Coutelis et al., 2013). Based on these data a model has been proposed in which *Abd-B* controls *myo1D* to direct dextral genitalia rotation, and a putative gene, negatively regulated by *myo1D*, to direct sinistral rotation (Coutelis et al., 2013; Coutelis et al, 2014; Geminard et al., 2014).

Genital plate rotation is defined in relation to a fixed coordinate element, the posterior abdomen, which serves as a reference frame, but which undergoes extensive remodeling during the same developmental period. In males and females, the adult abdominal epidermis is formed from histoblast nests, which proliferate during pupal development and are surrounded by polyploid larval epithelial cells (LECs), which undergo cell death and extrusion as the histoblast nests expand (Madhavan and Madhavan, 1980; Ninov et al., 2007). In addition, the male most posterior abdominal segment, the A7, is extruded under the control of *Abd-B* and the sex determination pathway (Wang et al., 2011; Foronda et al., 2012; Wang and Yoder, 2012). These abdominal events overlap temporally with genital disc rotation, and the disc physically contact posterior abdominal tissues during its movement, raising the question of a possible contribution of changes in abdominal epidermis development on genital disc circumrotation.

Here we address this question by studying how morphogenesis of the adult abdominal epidermis impacts genital disc rotation. We have found that circumrotation can proceed independently of male A7 elimination, and identify posterior abdominal LECs as the primary tissue contacting the rotating disc before and during rotation. We also discovered that the timely extrusion of LECs from the male abdominal epidermis, under the control of the cell death and the Epidermal Growth Factor (EGF) pathways, is essential for normal genital disc movement. We further show that timely clearance of Extracellular Matrix, mediated by metalloproteinases, is also needed for a complete rotation, and that *Abd-B* acts in both LECs and histoblasts. Together, our results reveal that genital disc circumrotation depends not only on an intrinsic Myo1D-driven program, but also on coordinated remodeling of adjacent posterior abdominal tissues that creates a permissive mechanical environment for rotation.

## Results

### The elimination of the male A7 is not required for complete genital disc rotation

Elimination of the male A7 and terminalia rotation are two Abd-B-dependent processes in the *Drosophila* male (Wang et al., 2011; Foronda et al., 2012; Coutelis et al., 2013). To analyze male A7 development we used the *MD761*-Gal4 strain, which drives expression in the A7 (Foronda et al., 2012). By using this line, we found that several mutant combinations, such as overexpression of the EGFR pathway or reduction of *extramacrochetae*, not only induce the formation of a small A7, but also disturbs the normal orientation of the terminalia (Fig. 1B-C’, D, compare with the control in A, A’, D). Internal dissection of male abdomens overexpressing EGFR showed dextral but incomplete rotation, indicating a defect in execution of circumrotation rather than a change in chirality (Fig. 1E; n=12). Mutations in *capicua*, which activate the EGFR pathway, have also been reported to affect relative position of male genitalia and analia (Forés et al., 2017).

**Figure 1.**
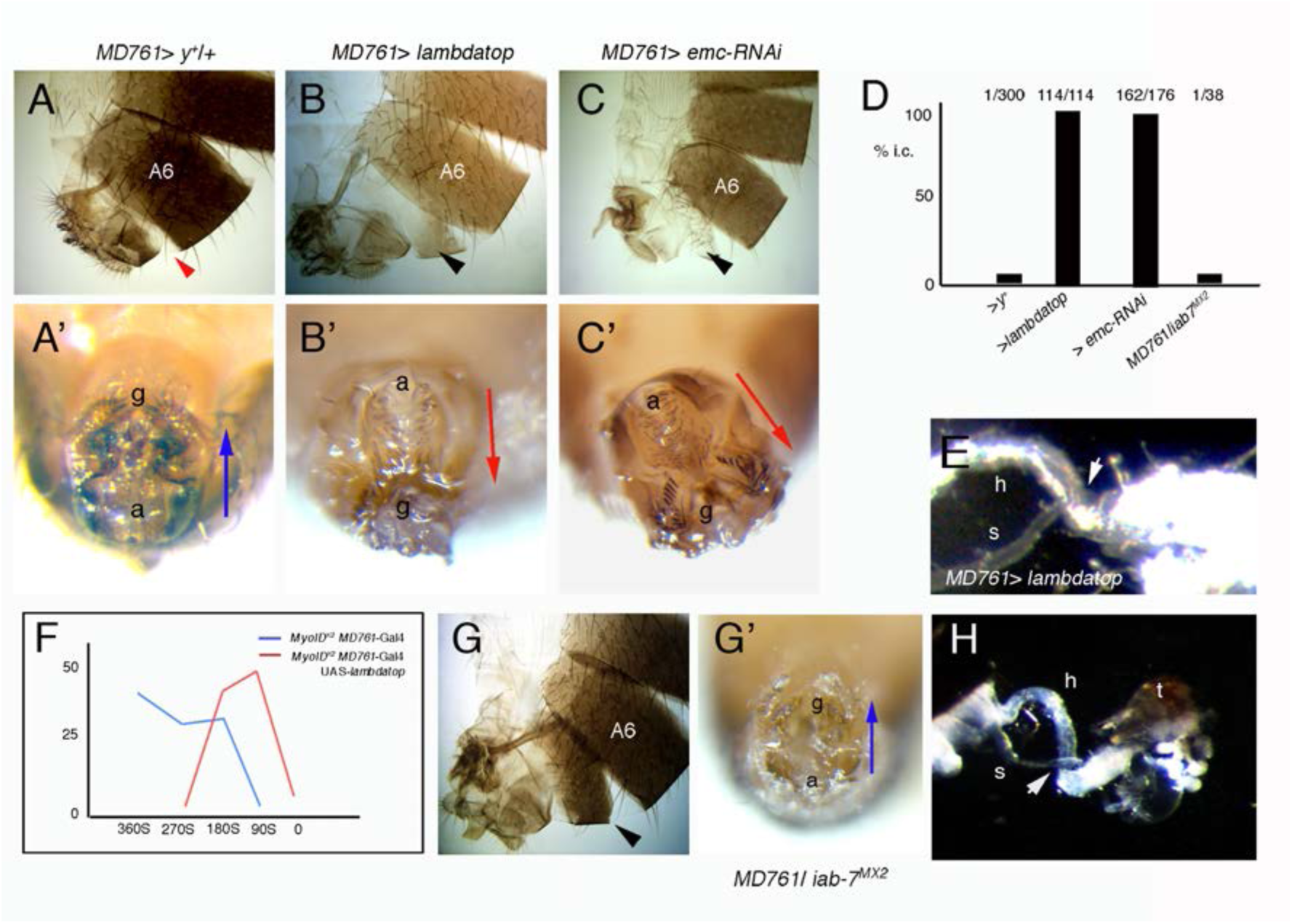
Suppression of male A7 extrusion is not required for normal genital plate rotation. A-C’) The expression of an activated form of the EGFR receptor (B, B’: *MD761*-Gal4 *tub*-Gal80^ts^ UAS-*Egfr^lambdatop^*; abbreviated in this and following Figures as lambdatop), or the knock down of *emc* (C, C’; *MD761*-Gal4 *tub*-Gal80^ts^ UAS-emcRNAi) in the male A7 induces the formation of a small A7 (B, C, black arrowhead) and abnormal genitalia (g) and analia (a) location (B’, C’) (compare with the *MD761*-Gal4 UAS-*y^+^* control in A, A’). D) Percentage of males with incomplete circumrotation (i. c.) of the terminalia in flies of the different genotypes (number of males with abnormal orientation divided by the total number of males studied are indicated). E) Looping of the spermiduct (s) on top of the hindgut (h) (arrow), in a *MD761*-Gal4 UAS-*lambdatop* male, indicating dextral rotation of the genital disc (Spéder et al., 2006). F) Graphical representation of the degree of sinistral rotation of males of the following genotypes: *myo1D^K2^*/*myo1D^K2^* (in blue; n=59) and *myo1D^K2^*/*myo1D^K2^ MD761*-Gal4 UAS-*Egfr^lambdatop^* (in red; n=65). Abscises shows the degrees of sinistral (S) rotation (from 0 to 360°S) and ordinates the percentage of males with a certain sinistral rotation. The activation of the EGFR pathway reduces the degree of sinistral rotation caused by the *myo1D^K2^* mutation as compared to the controls. G, G’) Posterior abdomen and terminalia of a *MD761*-Gal4/*iab-7^MX2^* male, showing a big A7 (G) but normal terminalia orientation (G’). A6, sixth abdominal segment; a, analia; g, genitalia; black arrowheads indicate presence of a small segment and red arrowheads, absence of an A7. Blue and red arrows go from analia (a) to genitalia (g) and indicate normal or abnormal genital disc rotation, respectively. H) Looping of the spermiduct (s) on top of the hindgut (h) (arrow), in a *MD761*-Gal4/*iab-7^MX2^* male, indicating dextral rotation of the genital disc; t, terminalia.

To test if the increase of EGFR activity affects directionality of rotation, we activated EGFR signaling in a Myosin 1D (Myo1D) mutant background, in which genital disc rotation is inverted (from dextral to sinistral) (Speder et al., 2006; Hozumi et al., 2006). EGFR activation reduced the extent of sinistral rotation rather than enhancing it (Fig. 1F; n=59-65), indicating that EGFR signaling limits circumrotation independently of the Myo1D-dependent chirality mechanism.

One possible interpretation of our data is that failure to eliminate the male A7 leads to abnormal genital plate rotation. However, we observed that mutations in the *infraabdominal-7* (*iab-7*) region of the bithorax complex, which directs the A7 expression of *Abd-B* (Karch et al., 1985; Celniker et al. 1990; Gyurkovics et al., 1990; Sánchez-Herrero, 1991), result in a large A7 (transformed into an anterior segment) but show normal arrangement of genitalia and analia (Fig. 1D, G G’). Dissection of these mutants reveals the full, dextral, 360° rotation (Fig. 1H) (n=12). Therefore, this result contradicted the assumption that elimination of the A7 is the cause of incomplete terminalia rotation and demonstrates genetic independence of the two processes.

### The rotating genital disc contacts posterior abdominal larval epidermal cells

Since A7 suppression was not required for genital disc rotation, and our results affecting this process were obtained with *MD761*-Gal4, we decided to examine the expression driven by this line in the posterior abdominal region with more detail. Close inspection of *MD761*-Gal4 UAS-GFP pupae of about 24-25h APF (when the genital plate starts its rotation) revealed a strong GFP signal in A7 (both in histoblasts and LECs), but also a weaker, but clear, expression in LECs posterior to this segment and just anterior to the genital disc (Fig. 2A). We define this region, which has just LECs and no histoblasts, as A8 abdominal segment. This is not to be mistaken for the genital A8, forming part of the genital disc (Nöthiger et al. 1977; Schüpbach et al., 1978).

**Figure 2.**
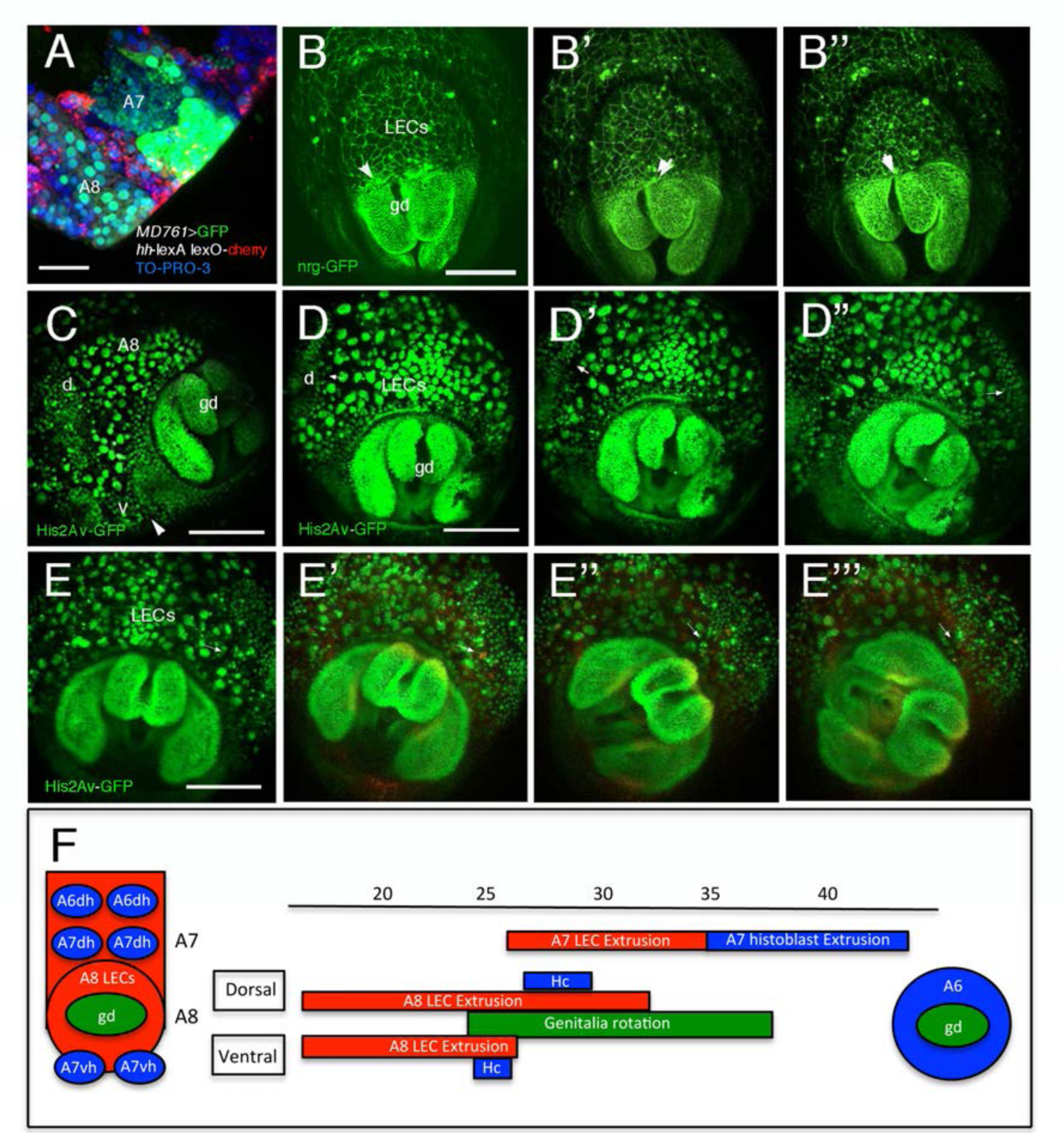
Genital plate rotation in relation with posterior abdomen development. A) Posterior abdominal epidermis of a UAS-GFP/+; *MD761*-Gal4 *hh*-lexA/LexO-Cherry pupa at approximately 24h APF. Note that *MD761*-Gal4 is driving GFP expression not only in A7 but also in the LECs of the A8, posterior to the A7p *hh-*expressing band. B-B’’) Snapshots from Supplementary Movie 1. The genotype of the pupa is Neuroglian-GFP and the movie goes approximately from 17 to 22h APF. It shows the genital disc (gd) and adjacent LECs. Arrows point to LECs that are being extruded. C) Just at the start of rotation, or shortly after, the ventral histoblast nest of the A7 (v) contacts the rotating genital disc (gd) (arrowhead). The dorsal nest (d) of the same segment is also indicated. D-D’’) Snapshots from Supplementary Movie 2. It is a His2Av-GFP pupa from approximately 24 to 26h APF. Genitalia start to rotate and the A7 dorsal histoblast nest (arrow) approaches the genital plate as the LECs are being extruded. E-E’’’) Snapshots from Supplementary Movie 3. A Hist2Av-GFP pupa from approximately 25 to 32h APF, showing the A7 histoblast nests (arrows) contacting the genital disc laterally, as the disc rotates up to near 180°. There are still some LECs (A7 LECs) around the dorsal midline at about 32h APF. F) Scheme summarizing the timing (hours APF) of dorsal and ventral contacts between dorsal (dh) and ventral (vh) histoblast nests and the genital disc (gd) and the extrusion of LECs. For A7 LECs we do not distinguish between ventral and dorsal cells. Hc, Histoblast first contact between histoblasts and genital disc. Scale bars are 100 μm in A, C, D, E, and 50 μm in B.

Live imaging indicated that the genital disc contacts posterior abdominal tissues before and during circumrotation. A8 LECs are adjacent to the genital disc and begin to extrude, without the presence of histoblasts (Foronda et al., 2012), before rotation of the disc (Fig. 2B-B’’; Supplementary Movie 1). Some ventral histoblasts contact the disc just as it starts to rotate (arrowhead in Fig. 2C), whereas dorsal ones approach the disc at about 25-26h APF, when the genital plate has not yet reached 90 degrees turn (Fig. 2D-D’’ and Supplementary Movie 2). As the genital plate continues its rotation, more LECs extrude and some histoblasts make contact with the genital plate (arrows in Fig. 2E-E’’’ and Supplementary Movie 3). A summary of the time course of these contacts is shown in Fig. 2F. We note, therefore, that the genital disc contacts LECs before and during most of its rotation, and that LECs, rather than histoblasts, constitute the major tissue interface for rotation.

To know if contact of A7 histoblasts with the genital disc is required, we impaired A7 histoblast proliferation by expressing, with the *MD761*-Gal4 line, UAS constructs expressing *dacapo* (*dap*), *fizzy-related* (*fzr*), or knocking down the expression of *cyclin dependent kinase 1* (*cdk1*), conditions known to strongly reduce cell proliferation (Stern et al., 1993; de Nooij et al., 1996; Sigrist and Lehner, 1997), including histoblast proliferation (Nakajima et al., 2011; Michel and Dahmann, 2020). Reduction of the female A7 size in these genotypes confirms that cell division is downregulated (Supplementary Fig. 1), whereas in males the genitalia and analia are positioned as in the wildtype (Fig. 3A-D). This suggests that the genital plate does not need to contact A7 histoblasts for its normal rotation. Live imaging in *MD761*-Gal4 *tub*-Gal80^ts^ UAS-*cdk1-RNAi* males confirms that, similarly to the control males (Fig. 3E-E’’’ and Supplementary Movie 4), in the mutant combination the genital disc rotates normally even though it does not contact the A7 histoblast nests throughout its circumrotation (Fig, 3F-F’’’ and Supplementary Movie 5). Our results indicate that cell contact between A7 histoblast nests and genital disc is not required for circumrotation, and point to the posterior LECs are the critical abdominal tissue contacting the genital disc.

**Figure 3.**
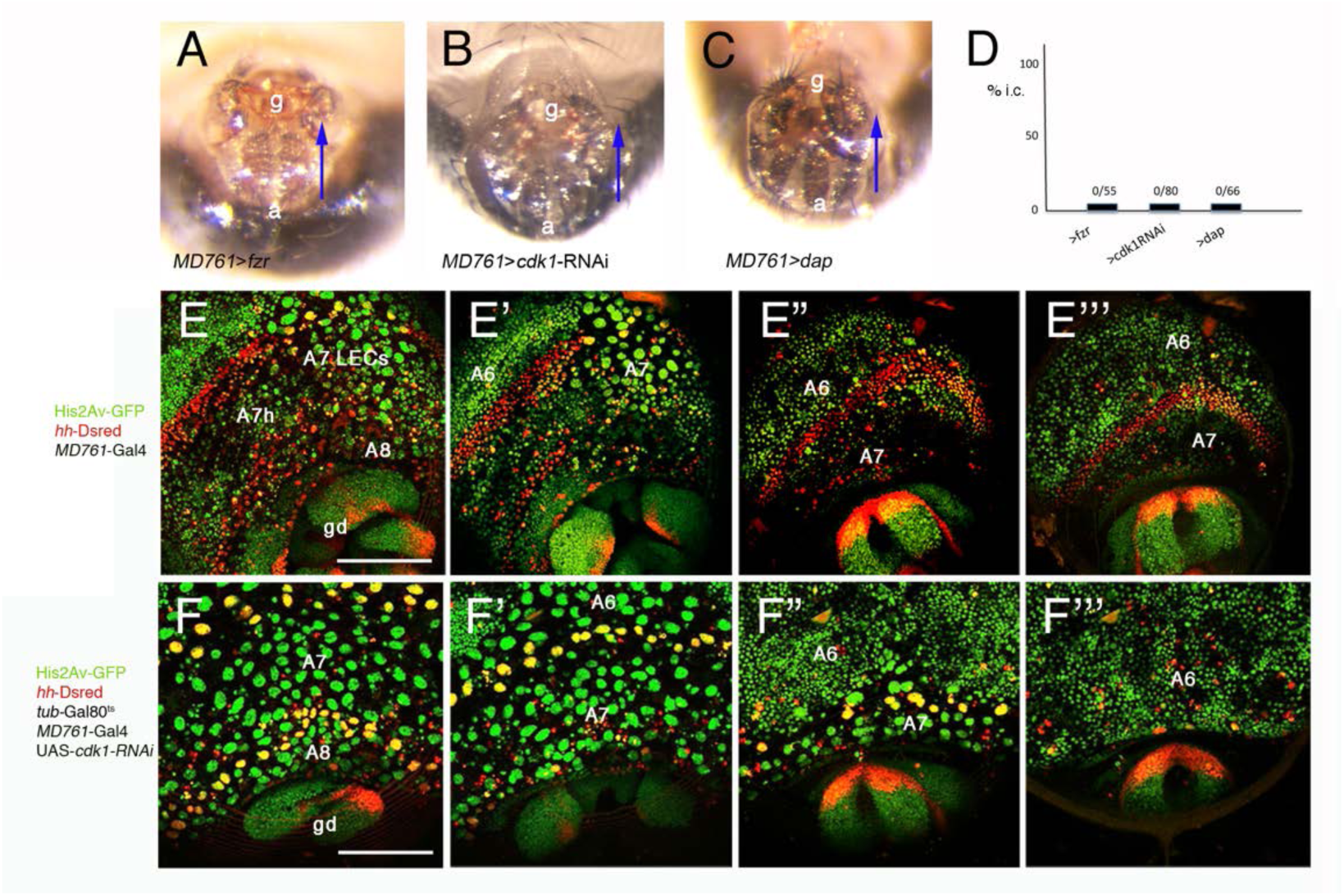
Reduction of cell division in male A7 histoblast nests does not prevent A8 LEC extrusion or genitalia rotation. A-C) Male genitalia and analia of *MD761*-Gal4 UAS-*fzr* (A), *MD761*-Gal4 UAS-cdk1-RNAi (B) and *MD761*-Gal4 UAS-*dap* (C) males (crosses made at 18°C and early third instar larvae shifted to 30°C). Symbols and arrows as in Fig. 1. D) Graphic showing percentage of incomplete circumrotation (i.c.) in these mutant combinations. Numbers and symbols as in Fig. 1. E-E’’’) Snapshots taken from Supplementary Movie 4, from approximately 27 to 37h APF, showing development of the posterior abdominal epidermis of a male of the genotype His2Av-GFP *MD761*-Gal4 *hh*-*DsRed*/+. Nuclei are marked by His2Av-GFP and posterior compartments are marked by *hh*-*DsRed*. When the genital disc (gd) ends its rotation, A8 LECs have already invaginated. F-F’’’) Snapshots taken from Supplementary Movie 5, from approximately 26 to 37h APF, showing development of the posterior abdomen of a male of the genotype UAS-cdk1-RNAi/*tub*-Gal80^ts^; His2Av-GFP/*MD761*-Gal4 *hh*-*DsRed*. Note that A8 LECs extrude and that A7 LECs extrude later than those in the A6 (and also later than the A7 LECs of the wildtype for a similar genital disc (gd) rotation time), suggesting that the absence of proliferation in histoblasts delays (but does not impede) LEC extrusion. Note also completion of genital disc rotation. g, genitalia; a, analia; A7h, A7 histoblasts. Scale bars are 100 μm in E, F.

### Apoptosis-driven extrusion of posterior abdominal LECs is required for complete circumrotation

In the pupal abdomen, LECs undergo cell death and extrude while histoblast nests proliferate and expand (Madhavan and Madhavan, 1980; Ninov et al., 2007; Bischoff and Cseresnyés, 2009; Nakajima et al., 2011; Kester and Nambu 2011; Baena-López et al., 2018; Michel and Dahmann, 2020). We detected cell death in LECs with *head involution defective* (*hid*)-EGFP-tagged (Nagarkar-Jaiswal et al., 2015) and UAS-Dronc-GFP-TETDG-Myc reporters (García-Arias et al., 2025), with a reporter of JNK activity, TRE-red (Chatterjee and Bohmann, 2012), or with Casexpress labeling (Ding et al., 2016; Tang et al., 2015) (Supplementary Fig. 2A-D’’; Supplementary Movie 6). Because posterior LECs contact the rotating genital disc, we tested if the cell death of abdominal LECs would also be a requisite for normal rotation.

When we prevented apoptosis in all LECs by using the LEC-specific *Eip71CD*-Gal4 line (Cherbas et al., 2003; Sekyrova et al., 2010; Singh and Mishra, 2014) to overexpress the mirRHG construct (Siegrist et al., 2010), or with *p35* (Hay et al., 1994), the genital disc did not complete circumrotation, as evidenced by terminalia orientation (Fig. 4B, C, G, the control in A). Similar results were obtained with the *MD761*-Gal4 line (Fig. 4E, F, G, the control in D). Examination of spermiduct looping in *Eip71CD*-Gal4 UAS-*mirRHG* males (n=4) show dextral incomplete rotation (Fig. 4H), indicating that the directional mechanism of rotation is intact when apoptosis is inhibited. Importantly, neither *MD761-*Gal4 (Fig. 4I, J; n=8) nor *Eip71CD*-Gal4 (Fig. 4K; n=6) drive expression in the genital disc, demonstrating that apoptosis inactivation just in LECs is sufficient to impair circumrotation. We conclude that apoptosis in posterior abdominal LECs is necessary for the correct movement of the genital plate.

**Figure 4.**
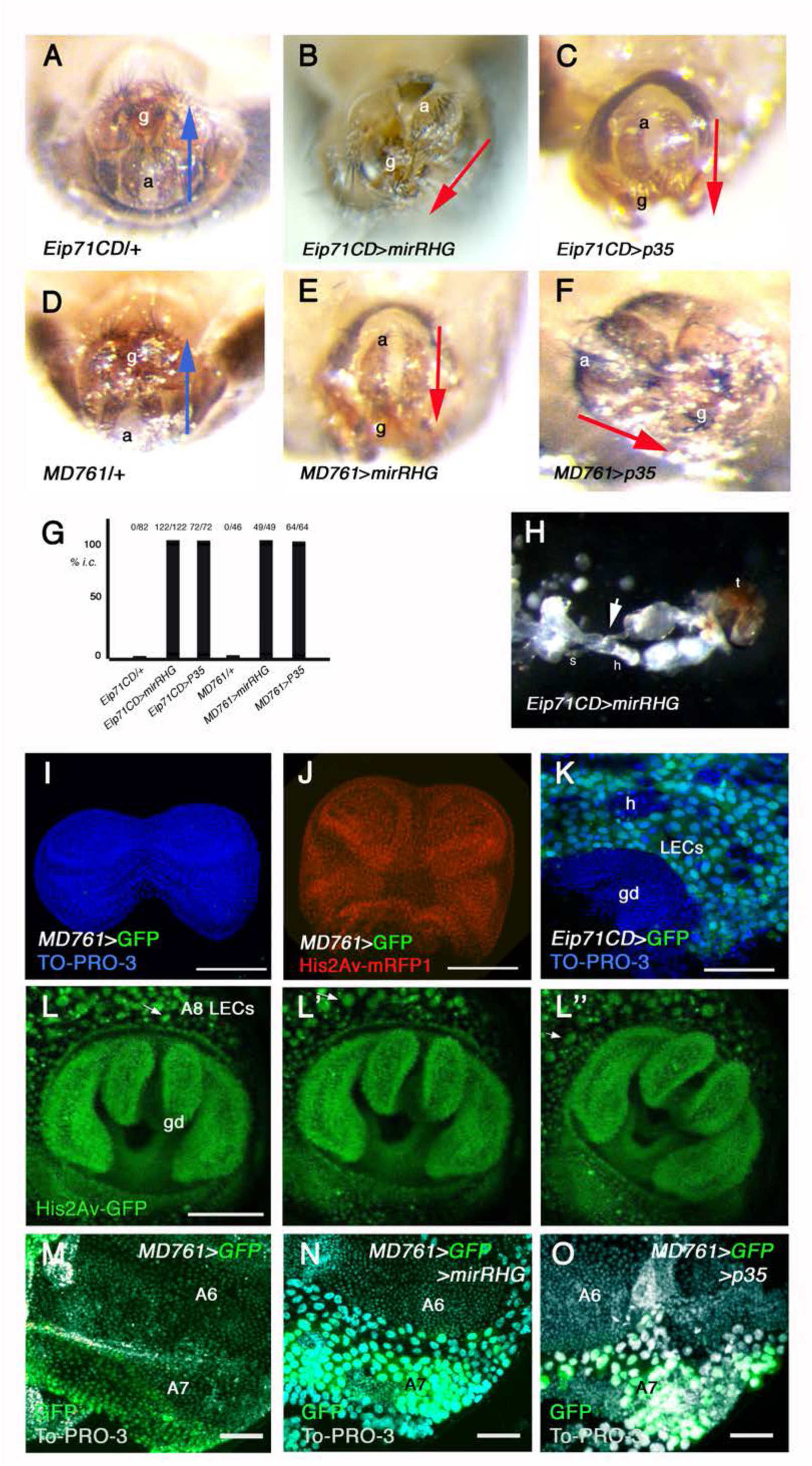
Cell death in the posterior LECs is required for a normal genital disc rotation. A, D) Normal position of genitalia and analia in *Eip71CD*-Gal4/+ (A) and *MD761*/+ (D) males. (B, C, E, F) Abnormal terminalia orientation in *Eip71CD*-Gal4 UAS-*mirRHG* (B), *Eip71CD*-Gal4 UAS-*p35* (C), *MD761*-Gal4 UAS-*mirRHG* (E) and *MD761*-Gal4 UAS-*p35* (F) males. G) Percentage of males with incomplete circumrotation (i.c.) in the four different genotypes, in which cell death is prevented. H) Internal dissection of the posterior part of a *Eip71CD*-Gal4 UAS-mirRHG male, showing normal spermiduct (s) on top of the hindgut (h) (arrow), indicating dextral turn of the genital disc; t, terminalia. I) Third instar male genital disc of genotype *MD761*-Gal4 UAS-GFP showing there is no GFP signal in the genital disc. J) Early pupa male genital disc of genotype *MD761*-Gal4 UAS-GFP His2Av-mRFP1 showing there is no GFP signal in the genital disc (His2Av-mRFP1 labels all nuclei). K) *Eip71CD*-Gal4 UAS-GFP posterior abdomen of a pupa of about 24h APF showing this Gal4 line drives expression in all the polyploid LECs but not in diploid histoblasts (h) or genital disc (gd) cells. TO-PRO-3 labels nuclei in blue. L-L’’) Snapshots from Supplementary Movie 7. His2AvGFP male pupa of about 26-29h APF showing extrusion of LECs adjacent to the genital disc (arrows) as it rotates. gd, genital disc. M-O) Posterior abdomens of pupae of about 38-40h APF of the genotypes *MD761*-Gal4 *tub*-Gal80^ts^ UAS-*GFP* (M), *MD761*-Gal4 *tub*-Gal80^ts^ UAS-*GFP* UAS-*mirRHG* (N), and *MD761*-Gal4 *tub*-Gal80^ts^ UAS-*GFP* UAS-*p35* (O). LECs have already extruded in the A6 in all the genotypes and in the A7 in M (control), but not in this A7 segment when cell death is inhibited (N, O). The genital disc was eliminated for easy mounting of the specimens and the A8 LECs are sometimes partially absent in the panels. M-O). The combination *MD761*-Gal4 *tub*-Gal80^ts^ is represented as *MD761^ts^*. 4-7 pupae were examined for each genotype in M-O. Blue and red arrows go from analia (a) to genitalia (g) and indicate normal or abnormal genital disc rotation, respectively. Scale Bars are 100 μm in I-L’’ and 50 μm in M-O.

Previous studies have shown that preventing apoptosis in LECs delays their extrusion (Ninov et al., 2007; Kester and Nambu, 2011; Nakajima et al., 2011; Teng et al., 2017; Michel and Dahmann, 2020). Detailed examination of movies (Fig. 2B-B’’; Fig. 4L-L’’; Supplementary movie 1; Supplementary Movie 7) reveal that LECs close to the genital disc are being extruded before and during genital disc rotation, suggesting a need of this timely extrusion for disc movement. We used the *MD761*-Gal4 line to examine cell behavior after selectively blocking apoptosis in posterior LECs. At about 35-40 h APF, the wild type A6 and A7/A8 LECs have already extruded (Fig. 4M). By contrast, inhibition of cell death by expressing the mirRHG construct or *p35* delays extrusion of the posterior LECs, while A6 LECs extrude as in the wildtype (Fig. 4N, O). Absence of LEC extrusion also result in an abdomen dorsal cleft phenotype (Ninov et al., 2007; Bischoff, 2012; Kester and Nambu, 2011) (Supplementary Figure 3A, B). Collectively, our results suggest that apoptosis is needed to drive extrusion of LECs contacting the genital disc for a correct circumrotation.

### Increased EGFR signaling disrupts circumrotation by delaying LEC elimination without changing chirality

Previous experiments have shown that increasing EGFR inhibits the activity of the proapoptotic genes *hid* and *reaper* (Bergmann et al., 1998; Kurada and White, 1998; Ewen-Campen and Perrimon, 2024), and prevents LECs elimination in pupae (Yuswan et al., 2024). Therefore, the effect on genital plate rotation observed after activation of the EGFR pathway (Fig. 1B’) is probably due to prevention or reduction of LEC cell death.

We tested the effect of activating the EGFR pathway with a constitutively active receptor (*Egfr^lambdatop^*; Queenan et al., 1997), a membrane-tethered Spitz ligand (*mSpi-GFP*; Schlesinger et al., 2004), or the *ras-1^V12^* mutant (Karim and Rubin 1998). Increasing Egfr signaling using either *MD761*-Gal4 or *Eip71CD*-Gal4 drivers consistently caused incomplete genital disc circumrotation (Fig. 1B’; 5A-G) while maintaining normal dextral rotation (Fig. 1E; Fig. 5H; n=5).

**Figure 5.**
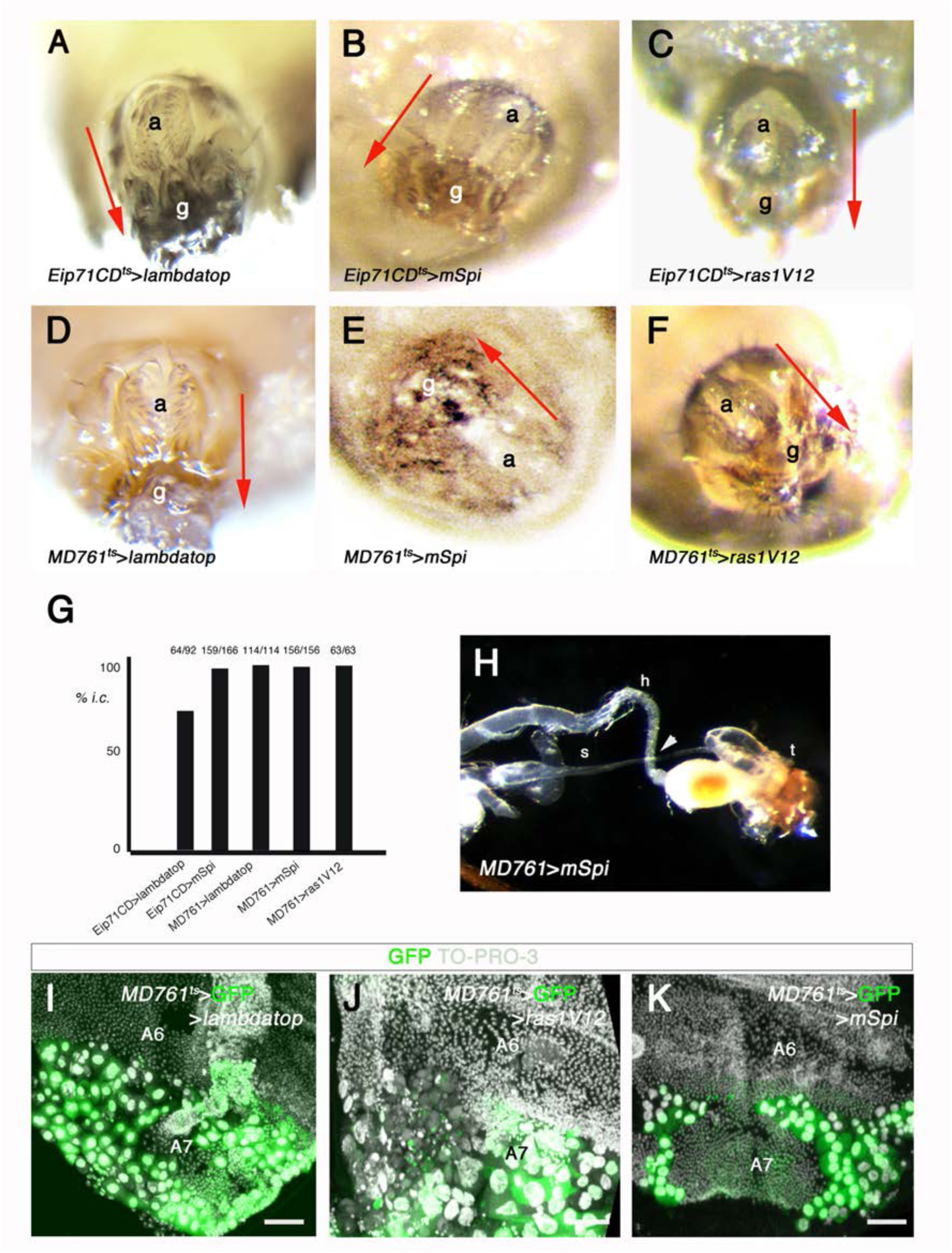
Normal levels of Egfr activity in the abdominal LECs are needed for a correct genital plate rotation. A) Abnormal orientation of genitalia and analia in males in which we express, with the *Eip71CD*-Gal4 *tub*-Gal80^ts^ (A-C) or the *MD761*-Gal4 *tub*-Gal80^ts^ (D-G) genetic combinations, UAS-*Egfr^lambdatop^* (A, D), UAS-*mSpi-GFP* (B, E), or UAS-*ras-1^V12^* (C, F). When *Eip71CD*-Gal4 UAS-*ras-1^v12^* late third instar larvae were shifted from 18°C to 30°C only three males survived, all of them having altered terminalia orientation. (G) Percentage of males with incomplete circumrotation (i.c.) in the different genotypes. (H) Internal dissection of a *MD761*-Gal4 UAS-*mSpi-GFP* male showing dextral movement of the genital disc (spermiduct, s, on top of hindgut, h; arrowhead; t, terminalia) (n=5). (I-K) Posterior abdomens of pupae of about 38-40h APF of the genotypes *MD761*-Gal4 *tub*-Gal80^ts^ UAS-*GFP* UAS-*Egfr^lambdatop^* (UAS-*lambdatop* in the panel) (I), *MD761*-Gal4 *tub-*Gal80^ts^ UAS-*GFP* UAS-*ras-1^V12^* (J) and *MD761*-Gal4 *tub*-Gal80^ts^ UAS-*GFP* UAS-*mSpi-GFP* (K): LECs have not been extruded in the posterior segments when the EGFR pathway is activated (compare with the A6 and with the control in Fig. 4M). The genital disc was eliminated for easy mounting of the specimens and the A8 LECs are sometimes partially absent in the panels I-K. The combinations *MD761*-Gal4 *tub*-Gal80^ts^ and *Eip71CD*-Gal4 *tub*-Gal80^ts^ are represented as *MD761^ts^* and *Eip71CD^ts^*, respectively. 4-7 pupae were examined for each genotype in N-P. Scale Bar is 50μm in I-K. Blue and red arrows go from analia (a) to genitalia (g) and indicate normal or abnormal genital disc rotation, respectively.

By activating the EGFR pathway with the *MD761*-Gal4 line, we found that posterior LEC extrusion was delayed relative to anterior segments (Fig. 5I-K, compare with 4M). As in the case of apoptosis inhibition, increasing EGFR activity also results in a failure of dorsal tergite fusion in adults, resulting in a dorsal cleft phenotype (Kester and Nambu, 2011; Singh and Mishra, 2015) (Supplementary Fig. 3C, D). Together with the results of spermiduct looping on top the hindgut (Fig. 1F), these results indicate that excessive EGFR signaling interferes with genital disc rotation by delaying apoptosis-driven LEC elimination, rather than by altering Myo1D-dependent chirality.

### Inhibition of Metalloproteinases also prevents genital disc circumrotation

The JNK pathway is active in LECs (Supplementary Fig, 2C, C’) and one of its targets, *Metalloproteinase 1* (*Mmp1*) (Uhlirova and Bohmann, 2006), is also present in these cells (Davis et al., 2022) (Fig. 6A; n=7). By contrast, we could not detect the expression of the other Metalloproteinase present in *Drosophila*, *Metalloproteinase 2* (*Mmp2*) (Fig. 6B) (n=8). We checked a possible role of Mmp1 in genital plate rotation by inhibiting its activity with the *Tissue inhibitor of Metalloproteinases* (*Timp*) gene. Expressing *Timp* with the *Eip71CD*-Gal4 or *MD761*-Gal4 drivers results in abnormal genitalia and analia orientation (Fig. 6C-E), while chirality is maintained (Fig. 6F) (n=8). However, although *Timp* expression phenocopies the effects of EGFR overactivation or apoptosis inhibition, the mechanism involved is different: the expression of *Timp* in A7/A8 does not delay extrusion of LECs with respect to the A6 (Fig. 6G) (n=6), and left and right hemitergites are normally fused at the midline if *Timp* is expressed in LECs (Supplementary Fig. 3E).

**Figure 6.**
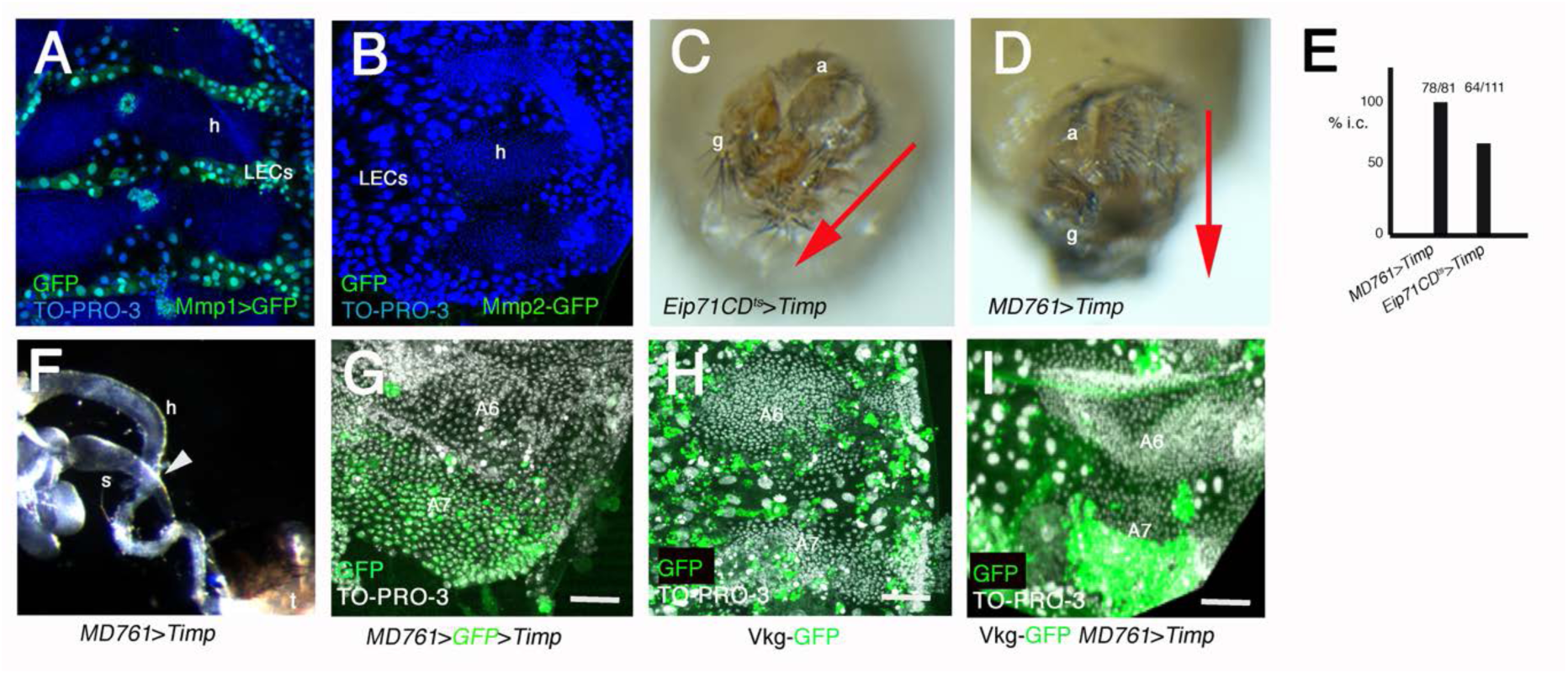
ECM clearance in the posterior abdominal epidermis is needed for genital plate circumrotation. A) Posterior abdomen of a pupa showing *Mmp1*-Gal4 UAS-GFP expression. Note strong signal in LECs but not in histoblasts (h). B) Mmp2-GFP pupal abdomen, showing no GFP signal. To-PRO-3 labels nuclei in blue in A and B. (C, D) Genitalia and analia of *Eip71CD-Gal4 tub*-Gal80^ts^ UAS-*Timp* (C) and *MD761*-Gal4 UAS-*Timp* males (D). See that reduction of Mmps activity in LECs prevents circumrotation of the genital plate. (E) Percentage of males with incomplete circumrotation (i.c.) in the two genotypes where *Timp* is expressed. F) Looping of the spermiduct (s) on top of the hindgut (h) in a *MD761*-Gal4 UAS-*Timp* male (arrowhead), showing dextral chirality (n=7). G) Posterior abdomens of pupae of about 36h APF of the *MD761*-Gal4 UAS-*GFP* UAS-*Timp* genotype, showing no effect on the timing of A7 LEC extrusion when inactivating metalloproteinases. (H, I) Posterior abdomens of about 24h APF pupae of the genotypes *vkg*-GFP (H) and *vkg*-GFP *MD761*-Gal4 UAS-*Timp* (I), showing accumulation of Vkg-GFP in the A7 when inactivating metalloproteinases. Scale bars are 100 μm in A, B and 50 μm in G-H. Blue and red arrows go from analia (a) to genitalia (g) and indicate normal or abnormal genital disc rotation, respectively.

The major components of the extracellular matrix (ECM) are progressively degraded in abdominal cells from 13h APF onwards, being low at 22-25h APF, when the genital plate starts to rotate (Davis et al., 2022). Using a collagen reporter, Viking-GFP (Vkg-GFP; Morin et al., 2001), we found a strong GFP signal in the posterior segments of Vkg-GFP *MD761*-Gal4 UAS-*Timp* abdomens (n=6), while the signal is low in more anterior segments or in the control abdomens (Fig. 6H, I) (n=6). These data suggest that timely ECM clearance in posterior abdominal LECs is required for circumrotation independently of LEC extrusion.

### *Abd-B* is required in either histoblasts or LECs cells for circumrotation

*Abd-B* is needed for A7 suppression in the abdomen (Sánchez-Herrero et al., 1985; Wang et al., 2011; Foronda et al., 2012; Singh and Mishra, 2014), and in the genital disc for circumrotation (Coutelis et al., 2013). To learn if it was also needed in posterior LECs for genital disc rotation, we expressed an *Abd-B*-RNAi construct with the *Eip71CD*-Gal4 line and found normal orientation of terminalia (Fig. 7A, B), no effect in abdominal development (Singh and Mishra, 2014), and normal chirality (Fig. 7C; n=7). However, Abd-B inactivation with the *MD761*-Gal4 line results in abnormal arrangement of terminalia (Fig. 7B, D). These animals also present an A7, indicating abnormal development of the posterior abdomen (Fig. 7E). To check if the effect on genital plate rotation was due exclusively to the activity on histoblasts, we crossed the UAS-*Abd-B*-RNAi line with a stock carrying the *MD761*-Gal4 line and the *332.3-Gal80* construct, which inhibits Gal4 expression in LECs (Fig. 7G, compare with 7F; Pasakarnis et al., 2016; Athilingam et al., 2022) (n=4). In contrast with the effect of *Abd-B* reduction in both histoblasts and LECs, in these animals the terminalia are normally positioned (Fig. 7B, H) and the A7 development is of intermediate size (Fig. 7I). These results indicate that knocking down *Abd-B* in posterior LECs and A7 histoblasts, but not just in one of these cell populations, results in abnormal terminalia arrangement. We noticed that this effect of reducing *Abd-B* is not due to a strong delay in LECs extrusion (Fig. 7J; compare with Fig. 4M-O and Fig. 5I-K) (n=5).

**Figure 7.**
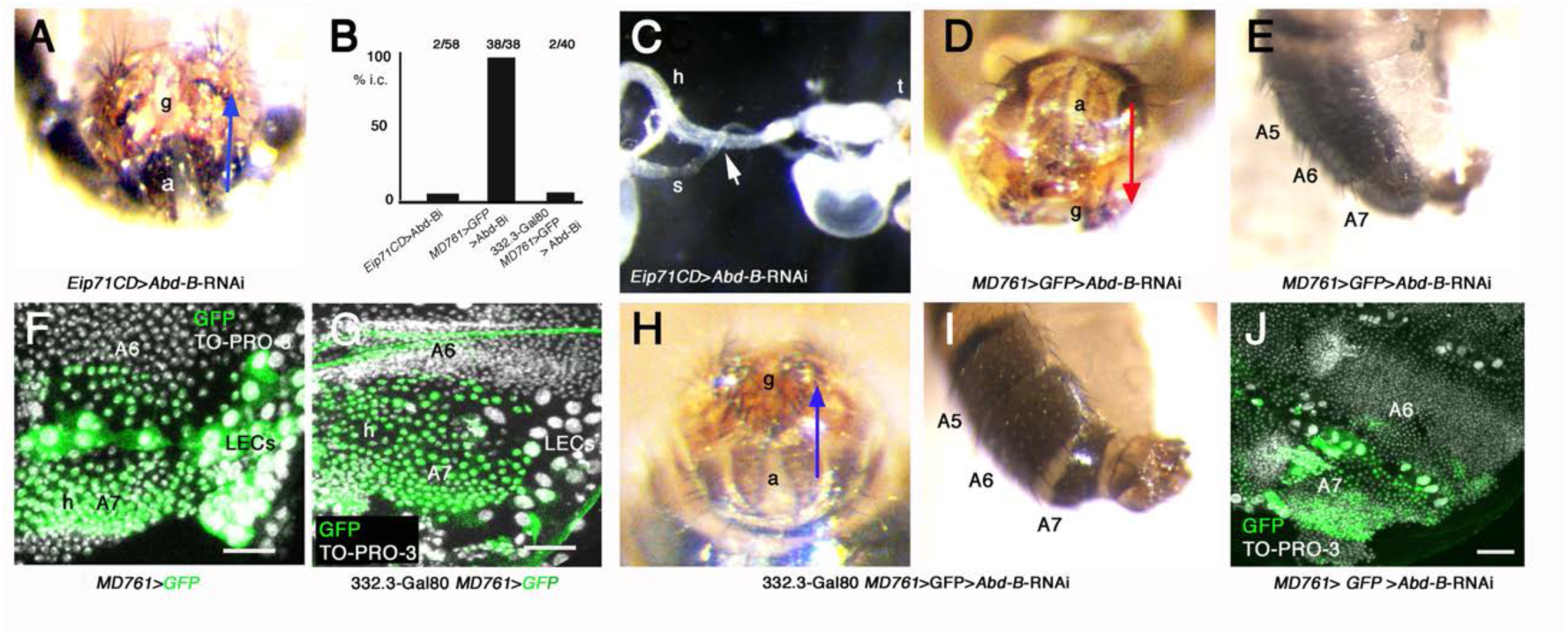
*Abd-B* activity in either histoblasts or LECs is sufficient for genital plate rotation. (A) The inactivation of *Abd-B* (*Eip71CD*-Gal4 UAS-Abd-B-RNAi in LECs does not modify the final orientation of the terminalia. B) Percentage of males with incomplete circumrotation (i.c.) in the different genotypes. C) Dissection of the posterior abdomen of a male showing the looping of the spermiduct (s) on top of the hindgut (h) in a *Eip71CD*-Gal4 UAS-Abd-BRNAi (arrow), showing dextral chirality; t, terminalia (n=7). (D, E) The inactivation of *Abd-B* in A7/A8 LECs and A7 histoblasts (*MD761*-Gal4 UAS-*GFP* UAS-Abd-BRNAi) changes the final location of genitalia and analia (D) and it also develops a well formed A7 (E). F) Posterior abdomen of an approximately 32h APF *MD761*-Gal4 UAS-*GFP* pupa, showing GFP expression in LECs and A7 histoblast nests. G) In the posterior abdomen of a 332.3-Gal80 *MD761*-Gal4 UAS-*GFP* pupa of a similar age there is expression of GFP just in A7 histoblasts. H, I) Genitalia and analia (H) and posterior abdomen (I) of 332.3-Gal80 *MD761*-Gal4 UAS-*GFP* UAS-Abd-BRNAi males, showing normal position of terminalia and development of an A7. 10-15 abdomens were studied for the genotypes shown in E and I. J) Pupa of about 38h APF. The down-regulation of *Abd-B* in posterior LECs and A7 histoblasts does not delay extrusion of LECs significantly; h, histoblasts. Scale bars are 50μm in G-I. Blue and red arrows go from analia (a) to genitalia (g) and indicate normal or abnormal genital disc rotation, respectively.

Together, these results show that genital disc circumrotation is independent of A7 elimination but depends on the combined Abd-B activity in posterior abdominal tissues.

## Discussion

The rotation of the *Drosophila* male genital disc depends on the intrinsic activity of Abd-B and Myo1D in the disc (Coutelis et al., 2013). Our results extend this view by demonstrating the requirement of posterior abdominal epidermal cells for this process. We have identified posterior LECs as a key extrinsic component and show that their timely elimination and the remodeling of ECM create a permissive mechanical environment for this rotation.

Pupal males undergo A7 segment elimination and 360° genital plate rotation. Both processes take place at the back of the pupa, present a short time overlap and depend on the Hox gene *Abdominal-B*. However, they are independent processes: the genital plate can undergo an incomplete rotation while a normal elimination of the A7 takes place, for example in *MD761* UAS-*p35* males (Foronda et al., 2012 and this report), and A7 extrusion can be prevented without affecting genital plate rotation. These observations demonstrate that A7 suppression is not required for circumrotation and clarify that previous correlations between the two processes do not reflect a strict functional dependency.

Our analysis points to posterior abdominal LECs as the critical tissue at the disc–abdomen interface. The genital disc contacts posterior LECs early and throughout most of its rotation, and these cells are being extruded as the movement proceeds. By contrast, histoblasts contact the genital disc when rotation has advanced and this contact is not needed, since blocking histoblast proliferation does not impair circumrotation (Fig. 4).

During pupal development, LECs die and extrude and histoblasts divide and occupy the space left to form the adult abdomen (Ninov et al., 2007; Bischoff and Cseresnyés, 2009; Nakajima et al., 2011; Michel and Dahmann, 2020). In LECs contacting the genital disc, this extrusion takes place before and during circumrotation. By removing cells that physically contact the rotating disc, apoptosis-driven extrusion likely reduces resistance at the disc–abdomen interface. Preventing apoptosis in posterior LECs, either directly or indirectly through elevated EGFR signaling, delays their extrusion and leads to incomplete rotation without altering chirality (Figs 4 and 5). These findings indicate that timely apoptosis-mediated LEC extrusion is required for the execution of circumrotation, but not for its directional specification, which remains controlled by Myo1D activity within the disc. Interestingly, apoptosis in the genital disc was also demonstrated to be needed for its rotation (Suzanne et al., 2010), indicating that cell death at adjacent tissues (genital disc, LECs) is a strict requirement for this movement.

We also found that remodeling of ECM is necessary for a normal genital disc rotation. *Mmp1* is specifically expressed in LECs of the abdomen and its activity promotes basal ECM degradation, which is largely completed by 21h APF (Davis et al., 2022), just before the start of genital plate rotation. Inhibition of metalloproteinase activity in posterior abdominal LECs results in persistence of collagen (Fig. 6), what also impairs circumrotation, but without delaying LEC extrusion. This suggests that ECM components contribute to the mechanical properties of the interface between the genital disc and the abdomen. Persistent ECM may increase stiffness or adhesion between genital disc and abdominal epidermis, opposing the forces that rotate the disc, as has been described in other cases: the stiffness of basement membrane in the Drosophila egg chamber, for example, exerts a compression that determines organ shape (Crest et al., 2017; Chiasta et al., 2017), and in *C. elegans*, the apical ECM prevents mechanical deformation by counteracting the pulling force of the elongating pharynx (Kelley et al., 2015).

We have shown that, apart from its role as controlling *myo1D* expression and genital disc direction of rotation, *Abd-B* is also needed in the posterior abdomen for circumrotation (Fig. 7). Downregulating *Abd-B* in either LECs or A7 histoblasts does not affect genital plate rotation, but the combined reduction in both tissues disturbs this movement. These observations indicate that, as to *Abd-B* requirement, posterior abdominal tissues contribute jointly to establish conditions permissive for circumrotation, and that disrupting only one tissue is insufficient to compromise this process.

Although we have shown independence of the male genital disc rotation and A7 extrusion, it is interesting to note that they seem to have evolved jointly in male Diptera. Primitive Diptera like mosquitoes have eight abdominal segments and their genital plate rotates 180°, while in more advanced Diptera there is a progressive reduction of male abdominal segments (6 in *Drosophila*, 5 in *Musca*) and a concomitant 360° rotation of the genital plate (Huber et al., 2007; Matsuda, 2013). These changes correspond with modifications of mating behaviors: in mosquitoes mating is of the end-to-end type, while in *Drosophila* it has evolved to the male-above position (Huber, 2010). This mating change, requiring 360° rotation of the genital plate, has been facilitated by reduction of the male abdomen number, thus coupling reduction of segment number, genital plate rotation and mating behavior (Yoder, 2012; Suzanne et al., 2010; Inatomi et al., 2019). This suggests that evolution may have acted on common regulatory modules to coordinate these male processes.

In conclusion, our study shows that genital disc circumrotation relies on the integration of intrinsic Myo1D-dependent forces within the disc and extrinsic remodeling of posterior abdominal tissues. This remodeling depends on the timely extrusion of LECs after apoptosis and on the elimination of ECM, thus removing constraints at the genital disc-abdominal epidermis interface and allowing disc rotation. These findings highlight the importance of tissue–tissue interactions and permissive mechanical environments in shaping complex morphogenetic movements.

## Material and Methods

### Drosophila Stocks

The *Drosophila melanogaster* wildtype line used is Vallecas. *iab-7*^MX2^ (Casanova et al., 1986) is a mutation in the *iab-7* regulatory region of *Abd-B*; *myo1D^k2^* is a mutation in *myo1D* (Spéder et al. 2006).

### Reporter lines

*His2*Av-GFP (BDSC 5941) and *His2*Av-mRFP1 (BDSC 23650) are histone H2A reporters. To label posterior compartments we used the insertion *hh*-*DsRed* (Kyoto DGRC 109137). (AP-1) TRE-DsRed identifies activity of the JNK pathway (BDSC 59011), *hid*-EGFP (BDSC 65331), *Mmp2*-GFP (Deady et al., 2015) and *vkgG454*-GFP (Morin et al., 2001; BDSC 98343) are insertions expressed as the corresponding genes*, and Nrg*-GFP is a fusion of GFP to the *Neuroglian* protein (Dubreuil et al., 1996).

### Gal4 and Gal80 lines

*MD761*-Gal4 (Foronda et al., 2012); *Eip71CD*-Gal4 (Cherbas et al. 2003) (BDSC 6871), *Mmp1*-Gal4 (BDSC 76161) and 332.3Gal80 (Pasakarnis et al., 2016). To detect caspase activation *in vivo* we used the Casexpress method (Ding et al. 2016; Tang et al. 2015).

### UAS lines

UAS-GFP (Ito et al., 1997), UAS-*p35* (Hay et al., 1994; BDSC 5072), UAS-*Egfr^lambdatop^* (BDSC 59843), UAS-*mSpi-GFP* (Tsruya et al. 2002), UAS-*ras-1*^V12^ (BDSC 64196), UAS-*Timp* (BDSC 58708), UAS-*y*^+^ (Calleja et al. 1996), UAS-*mirRHG* (Siegrist et al. 2010), UAS-*AbdB-*RNAi (VDRC 104872), UAS-*cdk1-RNAi* (BDSC 40950), UAS-*fzr* (Siles and Klämbt, 2010), UAS-*dap* (Lane et al., 1996), UAS-dronc-TETDG (García-Arias et al., 2005), UAS-*emc*-RNAi (Kyoto NIG-Fly 1007 R-2).

Temporal control of gene expression in the Gal4/UAS system was achieved with *tub*-Gal80^ts^ lines (McGuire et al., 2003) (BDSC 7017 and BDSC 7019). Larvae were normally shifted from 18°C to 30°C during third larval instar. *Eip71CD*-Gal4 *tub*-Gal80^ts^ UAS-*ras-1^V12^* larvae were shifted only at late third larval instar or white pupae stage due to the high mortality of shifting at earlier stages. Time at pupa was estimated according to the temperature of development. Some crosses were made at 30° (without *tub*-Gal80ts) to increase Gal4 activity.

### LexA/lexO system

*hh*-lexA (González-Mendez et al., 2017) and LexO-mCherry-CAAX (referred to in the manuscript as lexO-cherry; Yagi et al., 2010).

### Discs and pupal dissections

Timing of pupal development was set with respect to white pupae (0h after puparium formation, APF). To fix pupae they were placed on double-sided tape, dissected with a blade, washed with PBS to remove tracheae and fat body, fixed in 4% Paraformaldehyde (PFA) for 45 min and mounted in Vectashield (Vector laboratories, Inc.) for subsequent analysis, following established protocols (Foronda et al., 2012; Wang and Yoder, 2012). For the study of genital discs, larvae were dissected in cold PBS and fixation was carried out in 4% PFA for 30 min. Dissection, fixation and staining of imaginal discs were done according to standard protocols (Sullivan et al., 2000). TO-PRO-3 (Invitrogen) was used at a 1:1000 dilution to label nuclei.

### *In vivo* imaging

4D microscopy of pupae was performed as described in Seijo et al., 2015. Pupae were imaged with an inverted Leica SP8 or Zeiss LSM710 AxioObserver confocal microscope, the posterior end of the pupa facing down, and at 25°C. Pixel size was 512 x 512 pixels. Laser power was kept at the lowest possible level to avoid damaging the pupae. The images were further processed with Fiji / ImageJ (NIH) software (Schindelin et al., 2012). Time-lapse experiments were saved in video format at 6 frames per second for images taken every 10 min or more and at 15 frames for experiments where images were taken every 3 min. Genitalia movies were taken from the posterior of the pupa and are oriented dorsal up. At least 3 movies were obtained for each experiment.

### Determination of genital plate rotation chirality

In the cases indicated, we ascertained the genital disc rotation chirality through the dissection of the posterior abdomen of adults to determine the type of rotation of the spermiduct around the intestine (Spéder et al., 2006). The parallel observations of the position of the genitalia and the spermiduct rotation allow determining both the direction and the degree of rotation.

### Adult cuticle preparation

Flies were kept in a mixture of ethanol: glycerol (3:1) macerated in 10% KOH at 60 °C for 10 min, dissected, washed with water, dehydrated with ethanol and mounted in Euparal for inspection under a compound microscope.

### Image acquisition

Abdomens and genitalia that were not mounted were photographed with a M205FA dissecting scope with a DFC500 camera. Images from mounted adults were taken with a Zeiss Axiophot microscope. Confocal images were taken with the following microscopes: Leica TCS SPE, Leica TCS SP8, Zeiss LSM510, Zeiss LSM710, Zeiss LSM710 AxioObserver and Nikon A1R+ Eclipse Ti-E. Time-lapse experiments were carried out in Leica SP8 o Zeiss LSM710 AxioObserver inverted microscopes with a temperature control chamber at 25°C. Photographs of adult flies were taken with a Leica MZ12 stereomicroscope and a Leica DFC5000 camera, and images were acquired using Leica LAS software (3.7). The images were edited and assembled using Photoshop.

## Acknowledgments

We thank I. Guerrero for collaborating in some experiments and critical reading of the manuscript, L. A. Baena-López, K. Basler, D. Bohmann, D. Brunner, M. Calleja, C. Estella, I. Hariharan, D. Montell, G. Morata, B. Shilo, A. Spradling and S. Hayashi for stocks. We thank Mar Casado and Nuria Esteban for stock care and curation, the Confocal Microscopy Service at CBMSO, the Bloomington Stock Center, the Vienna Drosophila Resource Center, and the Kyoto Drosophila Stock Center for fly stocks. This study was supported by grants BFU2014-51989-P, BFU2017-86244-P and PID2020-113318GB-I00 funded by MICIU/AEI/10.13039/501100011033/and by ERDF A way of making Europe. N.P. was a recipient of a Formación del Personal Universitario (FPU) fellowship from the Spanish Government.

## References

1. Adám G, Perrimon N, Noselli S. The retinoic-like juvenile hormone controls the looping of left-right asymmetric organs in Drosophila. Development. 2003, 130, 2397–406. doi: 10.1242/dev.00460.

2. Athilingam T, Parihar SS, Bhattacharya R, Rizvi MS, Kumar A, Sinha P. Proximate larval epidermal cell layer generates forces for Pupal thorax closure in *Drosophila* Genetics. 2022 May 5;221(1): iyac030. doi: 10.1093/genetics/iyac030.

3. Baena-Lopez LA, Arthurton L, Bischoff M, Vincent JP, Alexandre C, McGregor R. Novel initiator caspase reporters uncover previously unknown features of caspase-activating cells. Development 2018 Dec 4;145(23):dev170811. doi: 10.1242/dev.170811.

4. Bergmann A, Agapite J, McCall K, Steller H. The Drosophila gene hid is a direct molecular target of Ras-dependent survival signaling. Cell. 1998 Oct 30;95(3):331–41. doi: 10.1016/s0092-8674(00)81765-1.

5. Bischoff M. Lamellipodia-based migrations of larval epithelial cells are required for normal closure of the adult epidermis of Drosophila. Dev Biol. 2012 Mar 1;363(1):179–90. doi: 10.1016/j.ydbio.2011.12.033.

6. Bischoff M, Cseresnyés Z. Cell rearrangements, cell divisions and cell death in a migrating epithelial sheet in the abdomen of Drosophila. Development. 2009; 136, 2403–11. doi: 10.1242/dev.035410.

7. Calleja M, Moreno E, Pelaz S, Morata G. Visualization of gene expression in living adult Drosophila. Science. 1996 Oct 11;274(5285):252–5. doi: 10.1126/science.274.5285.252.

8. Casanova J, Sánchez-Herrero E, Morata G. Identification and characterization of a parasegment specific regulatory element of the abdominal-B gene of Drosophila. Cell. 1986 Nov 21;47(4):627–36. doi: 10.1016/0092-8674(86)90627-6.

9. Celniker SE, Sharma S, Keelan DJ, Lewis EB. The molecular genetics of the bithorax complex of Drosophila: cis-regulation in the Abdominal-B domain. EMBO J 1990 Dec;9(13):4277–86. doi: 10.1002/j.1460-2075.1990.tb07876.x.

10. Chatterjee N, Bohmann D. A Versatile ΦC31 Based Reporter System for Measuring AP-1 and Nrf2 Signaling in Drosophila and in Tissue Culture. PLoS ONE 2012 7(4), e34063 doi: 10.1371/journal.pone.0034063.

11. Cherbas L., Hu X., Zhimulev I., Belyaeva E. y Cherbas P. EcR isoforms in Drosophila: testing tissue-specific requirements by targeted blockade and rescue. Development. 2003 Jan;130(2):271–84. doi: 10.1242/dev.00205.

12. Chlasta J, Milani P, Runel G, Duteyrat JL, Arias L, Lamiré LA, Boudaoud A, Grammont M. Variations in basement membrane mechanics are linked to epithelial morphogenesis. Development. 2017 Dec 1;144(23):4350–4362. doi: 10.1242/dev.152652.

13. Chougule A, Lapraz F, Földi I, Cerezo D, Mihály J, Noselli S. The Drosophila Actin Nucleator DAAM Is Essential for Left-Right Asymmetry. PLoS Genet. 2020 16(4): e1008758. doi: 10.1371/journal.pgen.1008758.

14. Coutelis JB, González-Morales N, Géminard C, Noselli S. Diversity and Convergence in the Mechanisms Establishing L/R Asymmetry in Metazoa. EMBO Rep 2014 Sep;15(9):926–37. doi: 10.15252/embr.201438972.

15. Coutelis JB, Géminard C, Spéder, P, Suzanne M, Petzoldt A, Noselli S. Drosophila left/right asymmetry establishment is controlled by the Hox gene abdominal-B. Dev. Cell. 2013; 24:89–97. doi: 10.1016/j.devcel.2012.11.013

16. Crest J, Diz-Muñoz A, Chen DY, Fletcher DA, Bilder D. Organ sculpting by patterned extracellular matrix stiffness. Elife. 2017 Jun 27;6: e24958. doi: 10.7554/eLife.24958.

17. Davis JR, Ainslie AP, Williamson JJ, Ferreira A, Torres-Sánchez A, Hoppe A, Mangione F, Smith MB, Martin-Blanco E, Salbreux G, Tapon N. ECM degradation in the Drosophila abdominal epidermis initiates tissue growth that ceases with rapid cell-cycle exit. Curr Biol 2022 Mar 28;32(6):1285–1300.e4. doi: 10.1016/j.cub.2022.01.045.

18. Deady LD, Shen W, Mosure SA, Spradling AC, Sun J. Matrix Metalloproteinase 2 Is Required for Ovulation and Corpus Luteum Formation in Drosophila. PLoS Genet 2015 Feb 19;11(2):e1004989. doi: 10.1371/journal.pgen.1004989.

19. de Nooij JC, Letendre MA, Hariharan IK. A cyclin-dependent kinase inhibitor, Dacapo, is necessary for timely exit from the cell cycle during Drosophila embryogenesis. Cell. 1996 Dec 27;87(7):1237–47. doi: 10.1016/s0092-8674(00)81819-x.

20. Ding A., Sun G, Argaw YG, Wong JO., Easwaran S, Montell DJ (2016). CasExpress reveals widespread and diverse patterns of cell survival of caspase-3 activation during development in vivo. Elife. 2016 Apr 8;5:e10936. doi: 10.7554/eLife.10936.

21. Dubreuil RR, MacVicar G., Dissanayake S, Liu C, Homer D, Hortsch M. (1996). Neuroglian-mediated cell adhesion induces assembly of the membrane skeleton at cell contact sites. J Cell Biol. 1996 May;133(3):647–55. doi: 10.1083/jcb.133.

22. Ewen-Campen B, Perrimon N. Wnt signaling modulates the response to DNA damage in the Drosophila wing imaginal disc by regulating the EGFR pathway. PLoS Biol. 2024 Jul 24;22(7): e3002547. doi: 10.1371/journal.pbio.3002547.

23. Forés M, Simón-Carrasco L, Ajuria L, Samper N, González-Crespo S, Drosten M, Barbacid M, Jiménez G. A new mode of DNA binding distinguishes Capicua from other HMG-box factors and explains its mutation patterns in cancer. PLoS Genet. 2017 Mar 9;13(3): e1006622. doi: 10.1371/journal.pgen.1006622.

24. Foronda D., Martín P. y Sánchez-Herrero E. Drosophila Hox and Sex-Determination Genes Control Segment Elimination through EGFR and extramacrochetae Activity. PLoS Genet. 2012; 8(8):e1002874. doi: 10.1371/journal.pgen.1002874.

25. García-Arias JM, Juárez-Uribe RA, Baena-López LA, Morata G and Sánchez-Herrero E. The Homeodomain-interacting protein kinase Hipk promotes apoptosis by stabilizing the active form of Dronc. Cell Death Discov. 2025 Dec 16;12(1):53. doi: 10.1038/s41420-025-02916-9.

26. Gleichauf R. Anatomie und Variabilität des Geschlechtsapparates von Drosophila melanogaster (Meigen) Z. Wiss. Zool. 1936; 148:1–66

27. González-Méndez L, Seijo-Barandiarán I, Guerrero I. Cytoneme-mediated cell-cell contacts for Hedgehog reception. Elife. 2017 Aug 21;6: e24045. doi: 10.7554/eLife.24045.

28. Gyurkovics H, Gausz J, Kummer J, Karch F. A new homeotic mutation in the Drosophila bithorax complex removes a boundary separating two domains of regulation. EMBO J. 1990 Aug;9(8):2579–85. doi: 10.1002/j.1460-2075.1990.tb07439.x.

29. Hay BA, Wolff T, Rubin GM. Expression of baculovirus P35 prevents cell death in Drosophila. Development. 1994 Aug;120(8):2121–9. doi: 10.1242/dev.120.8.2121.

30. Hozumi S., Maeda R., Taniguchi K., Kanai M., Shirakabe S., Sasamura T., Spéder P., Noselli S., Aigaki T., Murakami R. y Matsuno K. An unconventional myosin in Drosophila reverses the default handedness in visceral organs. Nature. 2006 Apr 6;440(7085):798–802. doi: 10.1038/nature04625.

31. Huber, B. A. Mating positions and the evolution of asymmetric insect genitalia. Genetica. 2010 Jan;138(1):19–25. doi: 10.1007/s10709-008-9339-6.

32. Huber BA, Sinclair BJ, Schmitt M. The evolution of asymmetric genitalia in spiders and insects. Biol Rev Camb Philos Soc. 2007 Nov;82(4):647–98. doi: 10.1111/j.1469-185X.2007.00029.x.

33. Inatomi M, Shin D, Lai YT, Matsuno K. Proper direction of male genitalia is prerequisite for copulation in *Drosophila*, implying cooperative evolution between genitalia rotation and mating behavior. Sci Rep. 2019 Jan 18;9(1):210. doi: 10.1038/s41598-018-36301-7.

34. Ito K, Awano W, Suzuki K, Hiromi Y, Yamamoto D. The Drosophila mushroom body is a quadruple structure of clonal units each of which contains a virtually identical set of neurones and glial cells. Development. 1997 Feb;124(4):761–71. doi: 10.1242/dev.124.4.761.

35. Karch F, Weiffenbach B, Peifer M, Bender W, Duncan I, Celniker S, Crosby M, Lewis EB. The abdominal region of the bithorax complex. Cell. 1985 Nov;43(1):81–96. doi: 10.1016/0092-8674(85)90014-5.

36. Karim FD, Rubin GM. Ectopic expression of activated Ras1 induces hyperplastic growth and increased cell death in Drosophila imaginal tissues. Development. 1998; 125:1–9. doi: 10.1242/dev.125.1.1.

37. Kelley M, Yochem J, Krieg M, Calixto A, Heiman MG, Kuzmanov A, Meli V, Chalfie M, Goodman MB, Shaham S, Frand A, Fay DS. FBN-1, a fibrillin-related protein, is required for resistance of the epidermis to mechanical deformation during C. elegans embryogenesis. Elife. 2015 Mar 23;4: e06565. doi: 10.7554/eLife.06565.

38. Kester RS, Nambu JR. Targeted expression of p35 reveals a role for caspases in formation of the adult abdominal cuticle in Drosophila. Int J Dev Biol. 2011;55(1):109–19. doi: 10.1387/ijdb.103109rk.

39. Kurada P, White K. Ras promotes cell survival in Drosophila by downregulating hid expression. Cell. 1998 Oct 30;95(3):319–29. doi: 10.1016/s0092-8674(00)81764-x.

40. Kuranaga E., Matsunuma T., Kanuka H., Takemoto K., Koto A., Kimura K., Miura M. Apoptosis controls the speed of looping morphogenesis in *Drosophila* male terminalia. Development. 2011 Apr;138(8):1493–9. doi: 10.1242/dev.058958.

41. Lane, ME, Sauer, K, Wallace, K. Jan, YN, Lehner, CF, Vaessin, H. Dacapo, a cyclin-dependent kinase inhibitor, stops cell proliferation during Drosophila development. Cell 1996. 87(7):1225–35. doi: 10.1016/s0092-8674(00)81818-8.

42. Madhavan MM, Madhavan K. Morphogenesis of the epidermis of adult abdomen of Drosophila. J Embryol Exp Morphol. 1980 Dec; 60:1–31.

43. Matsuda R. Morphology and Evolution of the Insect Abdomen: With Special Reference to Developmental Patterns and their Bearings upon Systematics. Pergamon Press 2013.

44. McGuire SE, Le PT, Osborn AJ, Matsumoto K, Davis RL. Spatiotemporal rescue of memory dysfunction in Drosophila. Science. 2003 Dec 5; 302(5651):1765–8. doi: 10.1126/science.1089035.

45. Michel M., Dahmann C. Tissue mechanical properties modulate cell extrusion in the *Drosophila* abdominal epidermis. Development. 2020 147(5). pii: dev179606. doi: 10.1242/dev.179606.

46. Morin X, Daneman R, Zavortink M, Chia W. A protein trap strategy to detect GFP-tagged proteins expressed from their endogenous loci in Drosophila. Proc Natl Acad Sci U S A. 2001 Dec 18;98(26):15050–5. doi: 10.1073/pnas.261408198.

47. Nagarkar-Jaiswal, S., Lee, P.T., Campbell, M.E., Chen, K., Anguiano-Zarate, S., Cantu Gutierrez, M., Busby, T., Lin, W.W., He, Y., Schulze, K.L., Booth, B.W., Evans-Holm, M., Nakajima Y, Kuranaga E, Sugimura K, Miyawaki A, Miura M. Nonautonomous apoptosis is triggered by local cell cycle progression during epithelial replacement in Drosophila. Mol Cell Biol. 2011 Jun;31(12):2499–512. doi: 10.1128/MCB.01046-10.

48. Ninov N, Chiarelli DA, Martin-Blanco E. Extrinsic and intrinsic mechanisms directing epithelial cell sheet replacement during Drosophila metamorphosis. Development. 2007 Jan;134(2):367–79. doi: 10.1242/dev.02728.

49. Nöthiger R, Dübendorfer A, Epper F. Gynandromorphs reveal two separate primordia for male and female genitalia in Drosophila melanogaster. Wilehm Roux Arch Dev Biol. 1977 Dec;181(4):367–373. doi: 10.1007/BF00848062.

50. Pasakarnis L, Frei E, Caussinus E, Affolter M, Brunner D. Amnioserosa cell constriction but not epidermal actin cable tension autonomously drives dorsal closure. Nat Cell Biol. 2016;18(11):1161–1172, doi: 10.1038/ncb3420.

51. Queenan AM, Ghabrial A, Schüpbach T. (1997). Ectopic activation of torpedo/Egfr, a Drosophila receptor tyrosine kinase, dorsalizes both the eggshell and the embryo. Development 1997 Oct;124(19):3871–80. doi: 10.1242/dev.124.19.3871.

52. Rousset R, Bono-Lauriol S, Gettings M, Suzanne M, Spéder P, Noselli S. The Drosophila serine protease homologue Scarface regulates JNK signalling in a negative-feedback loop during epithelial morphogenesis. Development. 2010; 37, 2177–86. doi: 10.1242/dev.050781.

53. Sánchez-Herrero E, Vernós I, Marco R, Morata G. Genetic organization of Drosophila bithorax complex. Nature. 1985 Jan 10-18;313(5998):108–13. doi: 10.1038/313108a0.

54. Sánchez-Herrero E. Control of the expression of the bithorax complex genes abdominal-A and Abdominal-B by cis-regulatory regions in Drosophila embryos. Development. 1991 Feb;111(2):437–49. doi: 10.1242/dev.111.2.437.

55. Sato K, Hiraiwa T, Maekawa E, Isomura A, Shibata T, Kuranaga E. Left-right Asymmetric Cell Intercalation Drives Directional Collective Cell Movement in Epithelial Morphogenesis. Nat Commun. 2015 Dec 10; 6:10074. doi: 10.1038/ncomms10074.

56. Schindelin J, Arganda-Carreras I, Frise E, Kaynig V, Longair M, Pietzsch T, Preibisch S, Rueden C, Saalfeld S, Schmid B, Tinevez JY, White DJ, Hartenstein V, Eliceiri K, Tomancak P, Cardona A. Fiji: an open-source platform for biological-image analysis. Nat Methods. 2012 Jun 28;9(7):676–82. doi: 10.1038/nmeth.2019.

57. Schlesinger A, Kiger A, Perrimon N, Shilo BZ. Small wing PLCgamma is required for ER retention of cleaved Spitz during eye development in Drosophila. Dev Cell. 2004; 7: 535–545. doi: 10.1016/j.devcel.2004.09.001.

58. Schüpbach T, Wieschaus E, Nöthiger R. The embryonic organization of the genital disc studied in genetic mosaics ofDrosophila melanogaster. Wilehm Roux Arch Dev Biol. 1978 Sep;185(3):249–270. doi: 10.1007/BF00848355.

59. Seijo I, Guerrero I, Bischoff M (2015). In vivo imaging of Hedgehog transport in Drosophila epithelia. Methods in Molecular Biology: Hedgehog Signalling Protocols. N. Riobo (ed.), Springer, Vol. 1322.

60. Sekyrova P, Bohmann D, Jindra M, Uhlirova M. Interaction between Drosophila bZIP proteins Atf3 and Jun prevents replacement of epithelial cells during metamorphosis. Development. 2010 Jan;137(1):141–50. doi: 10.1242/dev.037861.

61. Siegrist SE, Haque NS, Chen CH, Hay BA, Hariharan IK. Inactivation of Both Foxo and reaper Promotes Long-Term Adult Neurogenesis in Drosophila. Curr Biol. 2010 Apr 13;20(7):643–8. doi: 10.1016/j.cub.2010.01.060.

62. Sigrist SJ, Lehner CF. Drosophila fizzy-related down-regulates mitotic cyclins and is required for cell proliferation arrest and entry into endocycles Cell. 1997 Aug 22;90(4):671–81. doi: 10.1016/s0092-8674(00)80528-0.

63. Singh NP, Mishra RK. Role of abd-A and Abd-B in development of abdominal epithelia breaks posterior prevalence rule. PLoS Genet. 2014; 10, e1004717. doi: 10.1371/journal.pgen.1004717.

64. Singh NP, Mishra RK. Specific combinations of boundary element and Polycomb response element are required for the regulation of the Hox genes in Drosophila melanogaster. Mech Dev. 2015 Nov;138 Pt 2:141–150. doi: 10.1016/j.mod.2015.07.016.

65. Spéder P, Adám G, Noselli S. Type ID unconventional myosin controls left-right asymmetry in Drosophila. Nature 2006 Apr 6;440(7085):803–7. doi: 10.1038/nature04623.

66. Stern B, Ried G, Clegg NJ, Grigliatti TA, Lehner CF. Genetic analysis of the Drosophila cdc2 homolog. Development. 1993 Jan;117(1):219–32. doi: 10.1242/dev.117.1.219.

67. Sullivan W, Ashburner M, Hawley RS. (2000). Drosophila Protocols. Cold Spring Harbor Laboratory Press, New York.

68. Suzanne M., Pedtzoldt A.G., Spéder P., Coutelis J.B., Steller H. y Noselli S. Coupling of apoptosis and L/R patterning controls stepwise organ looping. Curr Biol. 2010 Oct 12;20(19):1773–8. doi: 10.1016/j.cub.2010.08.056.

69. Tang HL, Tang HM, Fung MC, Hardwick JM. In vivo CaspaseTracker biosensor system for detecting anastasis and non-apoptotic caspase activity. Sci Rep. 2015 Mar 11; 5:9015. doi: 10.1038/srep09015.

70. Teng X, Qin L, Le Borgne R, Toyama Y. Remodeling of adhesion and modulation of mechanical tensile forces during apoptosis in Drosophila epithelium. Development 2017; 144:95–105. doi: 10.1242/dev.139865.

71. Tsruya R, Sclesinger A, Reich A, Gabay L, Sapir A, Shilo B-Z. Intracellular trafficking by Star regulates cleavage of the Drosophila EGF receptor ligand Spitz. Genes Dev 2002 Jan 15;16(2):222–34. doi: 10.1101/gad.214202.

72. Uhlirova M, Bohmann D. JNK- and Fos-regulated Mmp1 expression cooperates with Ras to induce invasive tumors in *Drosophila*. EMBO J. 2006 Nov 15;25(22):5294–304. doi: 10.1038/sj.emboj.7601401.

73. Wang W., Kidd B.J., Carroll S.B. y Yoder JH. Sexually dimorphic regulation of the Wingless morphogen controls sex-specific segment number in Drosophila. Proc Natl Acad Sci U S A. 2011 Jul 5;108(27):11139–44. doi: 10.1073/pnas.1108431108.

74. Wang W, Yoder JH. Hox-mediated regulation of doublesex sculpts sex-specific abdomen morphology in Drosophila. Dev Dyn. 2012 Jun;241(6):1076–90. doi: 10.1002/dvdy.23791.

75. Yagi R, Mayer F, Basler K. Refined LexA transactivators and their use in combination with the Drosophila Gal4 system. Proc Natl Acad Sci U S A. 2010 Sep 14;107(37):16166–71. doi: 10.1073/pnas.1005957107.

76. Yoder JH. Fly (Austin). Abdominal segment reduction: development and evolution of a deeply fixed trait. 2012 Oct-Dec;6(4):240–5. doi: 10.4161/fly.22109.

77. Yuswan K, Sun X, Kuranaga E, Umetsu D (2024). Reduction of endocytosis and EGFR signaling is associated with the switch from isolated to clustered apoptosis during epitelial tissue remodeling in Drosophila. PLoS Biol. 2024 Oct 14;22(10):e3002823. doi: 10.1371/journal.pbio.3002823

