## Supplementary Figures for "*Drosophila* male genitalia rotation depends on permissive remodeling of the posterior abdomen"

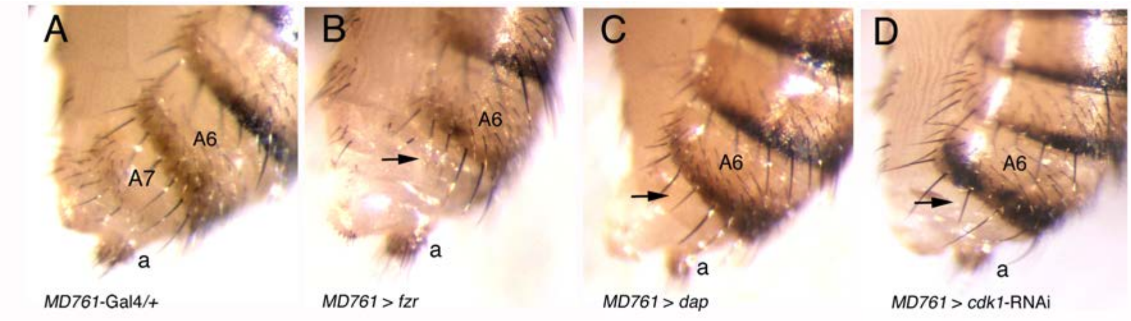


**Supplementary Figure 1. Reduction of female adult A7 segment when histoblast cell division is prevented.**

(A) Posterior abdominal epidermis of an *MD761*-Gal4/+ female. (B-D) Posterior female abdomen of flies of the following genotypes: *MD761*-Gal4 UAS-*fzr* (B), *MD761*-Gal4 UAS-*dap* (cross made at 31ºC) (C) and *MD761*-Gal4 UAS-*cdk1*-RNAi (D). Note the A7 segment is reduced in these genotypes (arrows). 15 females were checked for each genotype, all with a similar phenotype.

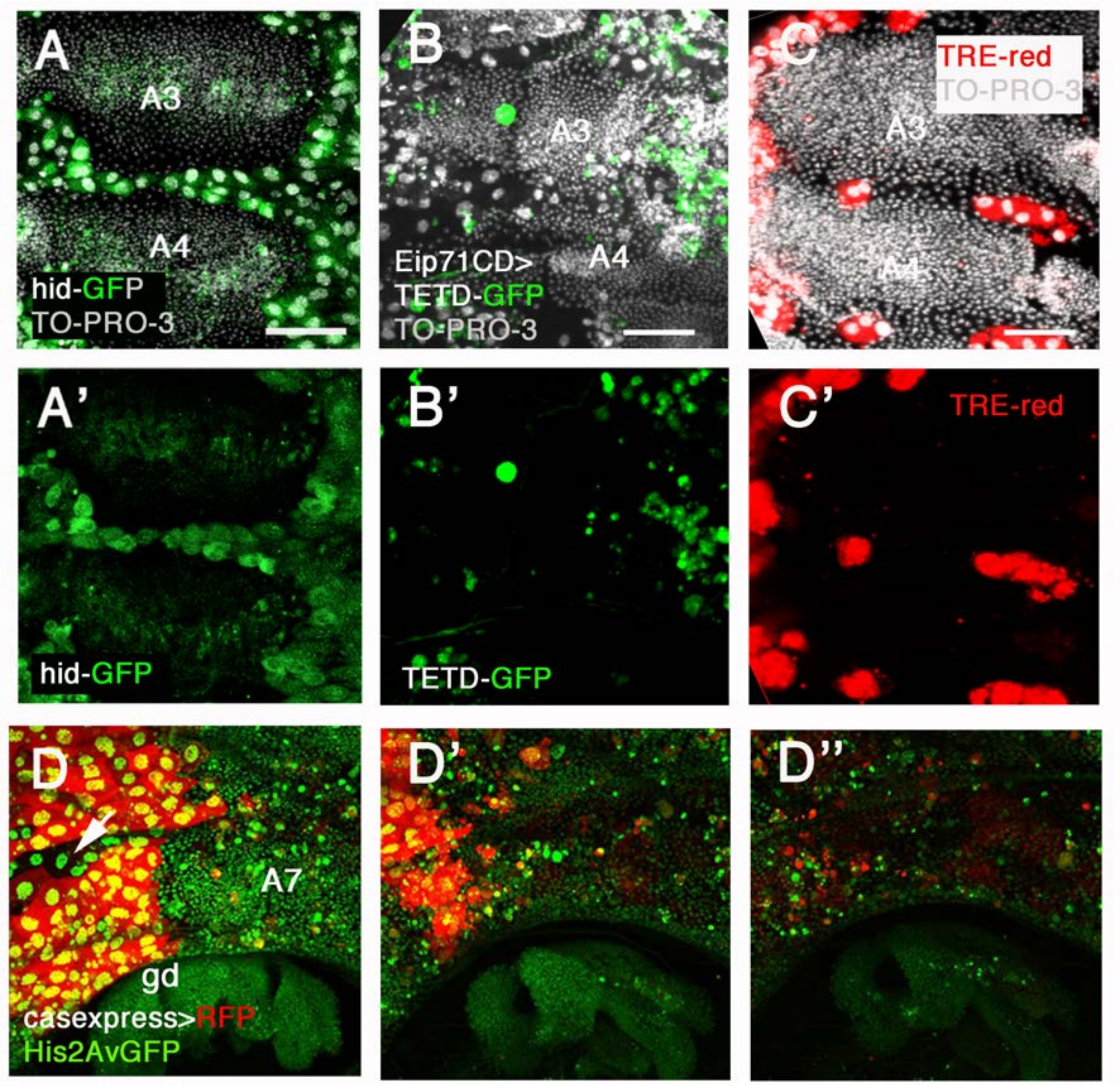


**Supplementary Figure 2. Cell death in the pupal abdomen**

(A-C') Two segments from abdomens of pupae of about 34-36h APF showing expression of *hid*-EGFP (A, A') in LECs and also in some histoblasts of the central region of the nest. This may indicate some unspecific signal of the construct, although Bischoff and Cseresnyés (Bischoff and Cseresnyés, 2009) report apoptosis in a few histoblasts in the wildtype (n=6). (B, B') *Eip71CD*-Gal4 UAS-TETDG-GFP abdomen presenting GFP signal in several LECs. The pattern of Dronc activation varies from abdomen to abdomen, which suggests that LECs are probably progressively dying and being extruded as development proceeds (n=5). (C, C') Abdomen of a TRE-red pupa. TRE marks most or all the LECs (n=7). D-D'') Snapshots from Supplementary Movie 7 showing the posterior abdomen of a pupa of about 35-37h APF of the genotype Casexpress UAS-RFP His2Av-GFP, showing the progressive elimination of LECs, marked in red. Note that a few LECs are not marked with RFP (arrow) but are equally extruded (see also Supplementary Movie 7). Scale bars are 100 μm.

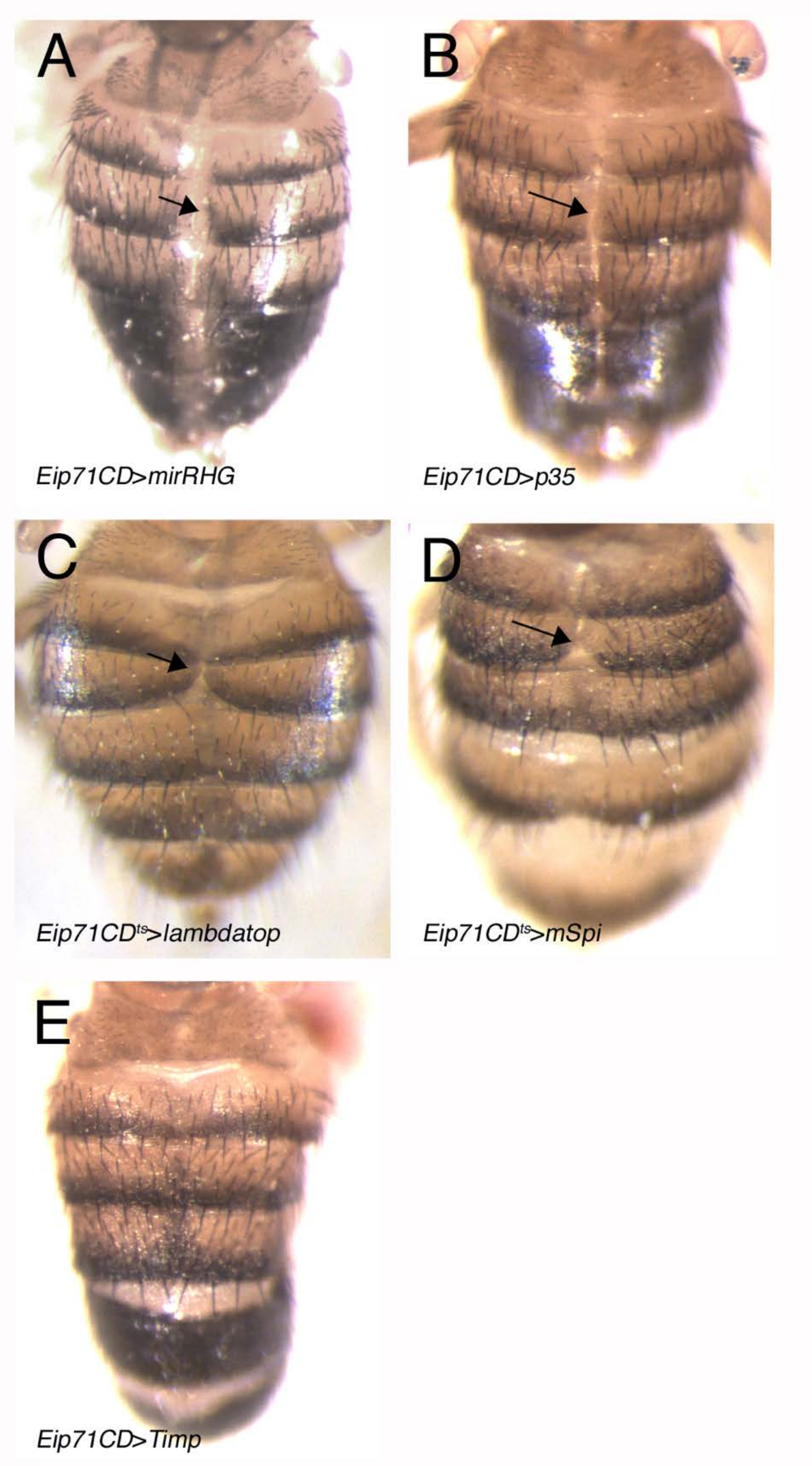

**Supplementary Figure 3. Effect of inactivating different genes or pathways in LECs on tergite fusion.**

Preventing cell death (*Eip71CD*-Gal4 UAS-*mirRHG*, A, or *Eip71CD*-Gal4, UAS-*p35*, B), or activating the EGFR pathway (*Eip71CD*-Gal4 *tub*-Gal80$^{ts}$ *Egfr*$^{lambdatop}$, *C,* or *Eip71CD*-Gal4 *tub*-Gal80$^{ts}$ *UAS-mSpi-GFP*, D) in LECs causes incomplete tergite fusion (arrows). However, expression of *Timp* (*Eip71CD*-Gal4 *tub*-Gal80$^{ts}$ UAS-*Timp*, E) in these cells does not result in tergite defects. C and D are females: the phenotypes are better observed in females than in males. 10-15 individuals were observed for each genotype, all of them showing some degree of tergite fusion defects.

**Supplementary Movie Legends**

**Supplementary Movie 1. Extrusion of LECs adjacent to the genital disc before it rotates.** The cells are marked by neuroglian-GFP (nrg-GFP). The arrows mark cells that are being extruded**.**

**Supplementary Movie 2. Start of genital plate rotation**. Hist2Av-GFP male, approx. 24h to 26h APF. Arrows mark dorsal A7 histoblast nests.

**Supplementary Movie 3. Contact of dorsal histoblast nests and genital disc**. Hist2Av-GFP male, approx. 25h-32h APF. Note how dorsal A7 histoblast nests approach and finally contact the genital plate (arrows).

**Supplementary Movie 4. Wildtype rotation, elimination of LECs and start of A7 histoblasts elimination.** Hist2Av-GFP *MD761*-Gal4 *hh-DsRed* male, approx. 27 to 37h APF. LECs are eliminated from A8 and A7; the A7 histoblast nests also begin to be extruded at the end of the movie, as the genital disc ends its rotation.

**Supplementary Movie 5. Suppression of male A7 histoblasts proliferation does not prevent correct genital plate rotation**. UAS-*cdk1*-RNAi/*tub*-Gal80$^{ts}$; His2Av-GFP *MD761*-Gal4 *hh-DsRed* male, approx. 26 to 37h APF. Note that LECs take longer to be extruded than in the control but finally do so. Note also the absence of the A7 histoblast nests, which do not proliferate (out of the field of view) and the contact of A6 histoblast nests with the genital plate, which rotates normally. Although A7 histoblasts do not proliferate, A7 LECs extrude.

**Supplementary Movie 6. The LECs show caspase activity and extrude.** Casexpress UAS-RFP Hist2Av-GFP male, approx. 30 to 42h APF. Note strong red label in almost all LECs and how they extrude as the genital disc rotates.

**Supplementary Movie 7. LECs adjacent to the genital plate extrude before its rotation.** Male of the His2Av-GFP genotype, approx. 26-29h APF. Note the extrusion of LECs of the A8 adjacent to the genital disc (a few of them are indicated by arrows).
